# Variance of allele balance calculated from low coverage sequencing data infers departure from a diploid state

**DOI:** 10.1101/2021.09.14.460322

**Authors:** Kyle Fletcher, Rongkui Han, Diederik Smilde, Richard Michelmore

## Abstract

**Motivation:** Polyploidy and heterokaryosis are common and consequential genetic phenomena that increase the number of haplotypes in an organism and complicate whole-genome sequence analysis. Allele balance has been used to infer polyploidy and heterokaryosis in diverse organisms using read sets sequenced to greater than 50x whole-genome coverage. However, Sequencing to adequate depth is costly if applied to multiple individuals or large genomes.

**Results:** We developed VCFvariance.pl to utilize the variance of allele balance to infer polyploidy and/or heterokaryosis at low sequence coverage. This analysis requires as little as 10x whole-genome coverage and reduces the allele balance profile down to a single value, which can be used to determine if an individual has two or more haplotypes. This approach was validated on simulated, synthetic, and authentic read sets from an oomycete, fungus, and plant. The approach was deployed to ascertain the genome status of multiple isolates of *Bremia lactucae* and *Phytophthora infestans*.

**Availability and implementation:** VCFvariance.pl is a Perl script available at https://github.com/kfletcher88/VCFvariance.

## Introduction

Polyploidy is a pervasive, well-known feature in all domains of eukaryotic life. Heterokaryosis is also common in eukaryotic microbes. Polyploidy is rarer in animals, but rife in plants where it has been described as a predominant component of sympatric speciation (Mable, 2004; Otto and Whitton, 2000). Increased cell size in plants is a consequence of polyploidy, possibly resulting in thicker, broader leaves and larger flowers, fruits, pollen, and stomata (Dar and Rehman, 2017; Otto and Whitton, 2000). Oomycetes (Stramenopiles), while somatically diploid, exhibit ploidy variation and/or heterokaryosis, which has consequences to fitness and virulence (Fletcher, et al., 2019; Knaus, et al., 2019). Many fungal species are somatically haploid, but like oomycetes, exhibit ploidy variation and/or heterokaryosis (Stajich, et al., 2009; Strom and Bushley, 2016; Zhu, et al., 2016). Flow cytometry can be used to estimate nuclear DNA content and can directly infer polyploidy when multiple genome sizes are found within a species (Bertier, et al., 2013; Catal, et al., 2010; Korpelainen, et al., 1997). However, this may not be technically feasible for some species or situations and is ineffective in detecting heterokaryosis.

High-throughput, whole-genome sequencing has enabled the detection of polyploidy and heterokaryosis in multiple species (Ament-Velásquez, et al., 2021; Bensasson, et al., 2019; Bertier, et al., 2013; Fletcher, et al., 2019; Fletcher, et al., 2018; Knaus, et al., 2019; Li, et al., 2016; Marburger, et al., 2018; Melo, et al., 2017; Strom and Bushley, 2016; Todd, et al., 2017; Tripp, et al., 2017; Weiß, et al., 2018; Yoshida, et al., 2013; Zhu, et al., 2016; Zhuang and Tripp, 2017). Several *in silico* approaches have been developed to summarize nucleotide frequencies at polymorphic sites and determine if two or more haplotypes exist in a sample (Knaus, et al., 2019; Weiß, et al., 2018). These approaches resolve the number of haplotypes present in a DNA sample by inferring the allele balance at bi-allelic single nucleotide polymorphisms (SNPs). A 0.5/0.5 balance indicates two haplotypes (likely diploid), 0.33/0.67 indicates three haplotypes (likely triploid), and 0.25/0.5/0.75 indicates four haplotypes (likely tetraploid). This approach was used to determine the ploidy of isolates of *Phytophthora infestans* (Li, et al., 2016; Yoshida, et al., 2013) and has since been deployed in plants, fungi, and animals as well as other oomycetes (Ament-Velásquez, et al., 2021; Bensasson, et al., 2019; Fletcher, et al., 2019; Fletcher, et al., 2018; Marburger, et al., 2018; Melo, et al., 2017; Tripp, et al., 2017; Weiß, et al., 2018; Zhu, et al., 2016; Zhuang and Tripp, 2017). In the case of the oomycete *Bremia lactucae*, several allele balance profiles were described for different isolates. Flow cytometry showed that the total DNA content of nuclei expected to be polyploid was the same as nuclei expected to be diploid. Upon further investigation, it was shown that heterokaryosis and not polyploidy was the cause of the observed variations in allele balance between isolates (Fletcher, et al., 2019). Generating allele balance histograms require adequate genome coverage to provide an unambiguous allele balance profile.

Sequencing to high depth (>50x) for adequate resolution is costly and a barrier to inferences using allele balance, especially when the genome is large. Analysis of individual SNPs in low-coverage data results in an allele balance profile that cannot resolve the number of haplotypes in an individual. Previously, a Gaussian Mixture Model was applied to classify allele balance into “diploid”, “triploid”, or “tetraploid” on downsampled datasets below 50x (Weiß, et al., 2018). This approach was reported to work well at lower coverage for *S. cerevisiae* (∼20x), but still required a higher read depth for analysis of *P. infestans* (Weiß, et al., 2018). Another approach that summarizes allele balance across multiple windows of a genome assembly was recently deployed in vcfR and was deployed on low coverage data (12x) to infer copy number variation of *Phytophthora infestans* isolates (Knaus and Grünwald, 2018; Knaus, et al., 2019). The latter method aims to determine if parts of the genome are present in higher quantities than expected and reports the proportions of inferred diploidy and polyploidy in each sample. This method requires subjective, specialist interpretation of its results.

The number of haplotypes in an individual will impact the sample variance of allele balance:

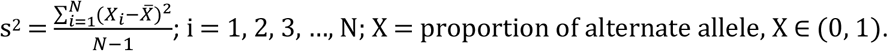

This is because in a diploid, the allele frequency of most SNPs will be 0.5/0.5, resulting in lower variance of the dataset. In a polyploid, many SNPs will not have an allele balance of 0.5/0.5, therefore increasing the variance in the dataset. Variance of allele balance has not previously been reported as an objective approach to discriminate between polyploids from diploids. In this paper, we establish a protocol for utilizing the variance of allele balance to detect departures from the diploid state. This protocol was validated on simulated, synthesized, and downsampled genuine high-coverage whole-genome sequencing data, from a plant, a fungus, and two oomycetes. The protocol was deployed to analyze 21 European race type isolates of *B. lactucae*, demonstrating that 12 were homokaryotic; the other nine were likely heterokaryotic. This protocol allows the use of existing low coverage datasets and generation of additional informative datasets at low cost.

## Methods

### Protocol validation

Simulated datasets were generated to capture allele frequencies for 100,000 loci with ploidy levels ranging from 2n to 8n at coverages ranging from 10x to 100x using rbinom() in R (R Development Core Team, 2012). Allele frequency profiles were filtered to only consider frequencies between 0.2 and 0.8, a common criterion to filter for heterozygous variants (Fletcher, et al., 2019; Li, et al., 2016; Yoshida, et al., 2013). Results were rounded to two decimal places, tabulated, and plotted using the base barplot(). Variance for each simulation was calculated with var() and plotted as a line graph using ggplot2 (Wickham, 2016). Variance calculations were replicated 1,000 times using repeat() to obtain mean and standard deviation statistics.

Diploid and polyploid genotypes were synthesized *in silico* from *Escherichia coli* strain K-12 (GCA_ 000005845.2) by introducing mutations into the reference assembly and proliferating them to the desired ploidy level. The number of polymorphisms at each ploidy level are detailed in Supplementary Table 1. Synthetic genotypes were generated using BedTools v2.25.0 maskfasta (Quinlan, 2014), masking with the nucleotide to be mutated. Synthetic genotype sequences were combined with the original assembly to generate templates for read generation. Synthetic 150 bp reads were generated from these combined assemblies using RandomReads.sh (Bushnell, 2016), generating 100x coverage.

Authentic reads of 30 *Bremia lactucae* isolates sequenced to high coverage and previously determined to be homokaryotic or heterokaryotic (Fletcher, et al., 2019) were downloaded from SRA BioProject PRJNA387192 (Supplementary Table 2). Reads of 40 *Phytophthora infestans* isolates sequenced to varying coverages were downloaded from several SRA accessions (Supplementary Table 3). These reads have been used previously to infer as diploidy, polyploidy or continuous copy number variation (Knaus, et al., 2019; Yoshida, et al., 2013). Reads of 24 *Saccharomyces cerevisiae* isolates that exhibited ploidy variation (Zhu, et al., 2016) were downloaded from SRA BioProject PRJNA315044 (Supplementary Table 4). Reads of 24 *Arabidopsis arenosa* accessions from 17 European populations, which were characterized as diploid or tetraploid (Monnahan, et al., 2019), were downloaded from SRA BioProject PRJNA484107 (Supplementary Table 5).

Simulation and genuine reads were deduplicated with SuperDeduper (Petersen, et al., 2015), then aligned to their respective GenBank reference sequences (*E. coli*; GCA_ 000005845.2, *B. lactucae*; GCA_004359215.1, *P. infestans*; GCA_000142945.1, *S. cerevisiae*; GCA_000146045.2, and *Arabidopsis arenosa*; GCA_905216605.1) using BWA-MEM v0.7.16 (Li, 2013). Variants were called with FreeBayes v1.2 (Garrison and Marth, 2012). The genome wide coverage was calculated by BEDtools (Quinlan, 2014). A Perl script (VCFvariance.pl) was implemented to calculate the variance of allele balance of SNPs filtered to be within 40% coverage of the genome wide coverage; this can be altered to include fewer or more polymorphisms with the -d flag. Allele balance between 0.2 and 0.8, mapping quality >30, and genotype quality >30 was filtered for to eliminate sequencing error. In addition, the Perl script flagged potential haploid accessions (e.g. in *S. cerevisiae*) if less than 55% of the SNPs passing coverage, mapping quality, and genotype quality filters failed the allele balance filter. This final option can be modified using the -p flag if necessary. For *E. coli* (synthetic polyploids), *B. lactucae*, and *S. cerevisiae*, BAM files were downsampled using SAMtools (Li, et al., 2009), and variant calling and variance calculation were repeated as above. Bar plots were generated with R base barplot() and table() functions (R Development Core Team, 2012). Variance of allele balance were plotted per individual using either geom_line() or geom_point() (Wickham, 2016) and labelled with ggrepel() (Slowikowski, 2017). For *B. lactucae*, kernel density estimation was plotted using geom_density_2d() (Wickham, 2016). Contours produced from the estimation of *B. lactucae* kernel density were also plotted in the *P. infestans, A. arenosa*, and *S. cerevisiae* figures for comparative analysis. VCFvariance.pl is available at https://github.com/kfletcher88/VCFvariance.

### Application of protocol

Low coverage sequencing of European type isolates of *B. lactucae* was used to demonstrate utility of this protocol. Spore pellets in ethanol for type isolates of European races of *B. lactucae* were provided by Diederik Smilde (Naktuinbouw, The Netherlands). DNA was extracted as described previously (Fletcher, et al., 2019). Paired-end (∼300 bp fragments) libraries were prepared using the Kapa HyperPrep kit following the manufacturer’s instructions (Roche, Switzerland). Libraries were sequenced on a MiSeq 2500 or HiSeq 4000. Reads were uploaded to NCBI BioProject PRJNA387192. Reads were deduplicated using SuperDeduper (Petersen, et al., 2015), trimmed for adapters and low quality sequence using BBMAP bbduk.sh (Bushnell, 2016) and mapped with BWA mem v0.7.16 (Li, 2013). SNPs were called with FreeBayes v1.2 (Garrison and Marth, 2012) and the variance of allele balance for each isolate was calculated using VCFvariance.pl (https://github.com/kfletcher88/VCFvariance). One point per isolate was plotted as coverage by variance. Kernel density estimates of established homokaryotic and heterokaryotic *B. lactucae* isolates were plotted as a guide using geom_density2d with six bins (Wickham, 2016). This plot was inspected to determine whether the isolates fit the model of a homokaryotic diploid.

## Results

The simulation test demonstrated that the variance of allele balance can discriminate between diploids and polyploids. Plotting histograms of simulated data recreated the expected allele frequency plots for ploidies 2n to 8n, provided over 50x coverage was simulated per site (Figure 1a, Supplementary Figure 1). As the coverage per locus declined, the resolution of the allele frequency plots decreased, although it was still possible to discriminate between diploids and polyploids in this model dataset at coverage as low as 10x (Supplementary Figure 1). Calculating the variance of allele balance demonstrated that there was a large difference between diploids and polyploids at all coverage levels ≥10x (Figure 1B). At 10x, the mean variance of allele frequencies for simulated diploids (2n) was 0.022 (+/- 8.46×10^−05^, 1,000 replicates), while simulated polyploids (3n to 8n) had variance of allele frequencies ranging from 0.035 to 0.40 (+/- 1.2×10^−04^, 1,000 replicates). As the coverage increased, the variance of allele balance for the simulated diploids declined; for simulated polyploids, the variance of allele balance remained constant (Figure 1B; Supplementary Table 6). The standard deviation of 1,000 replicates demonstrated that it was not possible to reliably decipher between different polyploids (>2n; Supplementary Table 6). Therefore, the difference between the variance of allele balance for diploids vs. polyploids means this metric can be used to discriminate diploids from polyploids.

**Figure 1.**
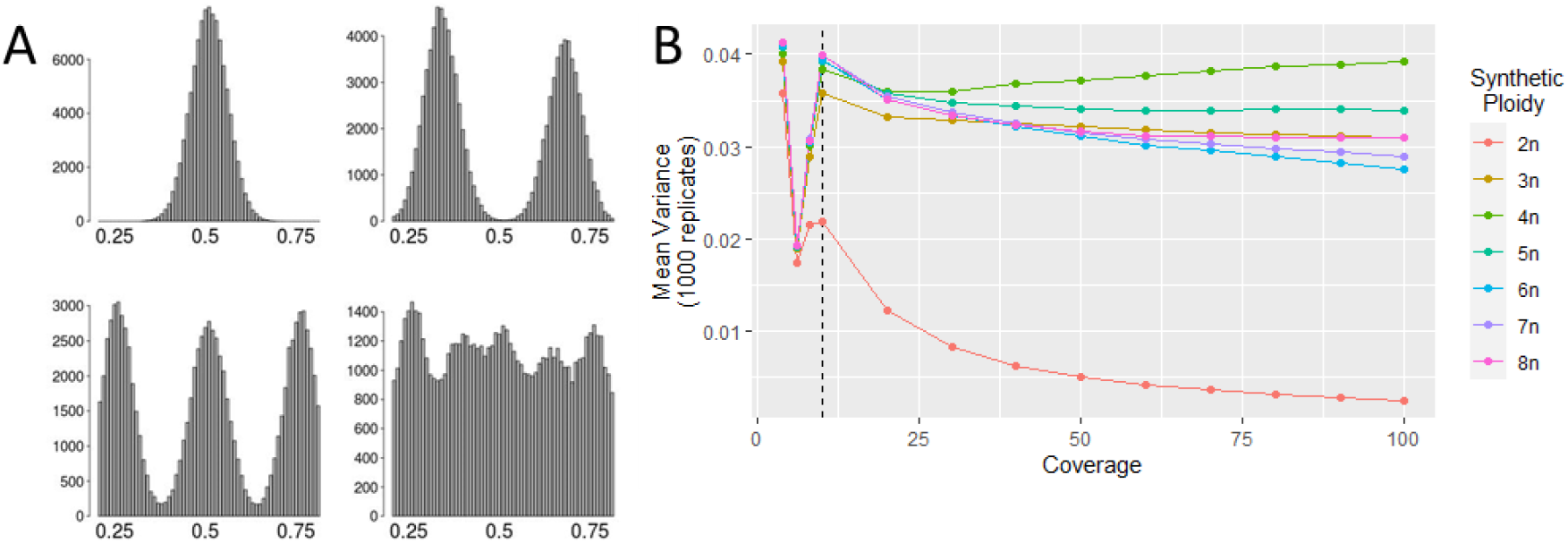
Statistical simulation of variance of allele balance at different ploidy levels. A) Allele balance histograms of 100,000 loci derived from 2n, 3n, 4n, and 8n datasets simulated to 100x whole-genome coverage. Downsampled allele balance histograms are presented in Supplementary Figure 1. B) Variance of allele balance calculated for 2n to 8n ploidies, with coverages at 6x, 8x, and 10x to 100x in increments of 10. The simulated diploid can be discriminated from simulated polyploids at coverages over 10x (dashed line). Below 10x, variance of allele frequency begins to converge for all ploidies.

Variance of allele balance was able to discern synthetic diploids from polyploids, generated from the *E. coli* genome sequence. Synthetic reads were analyzed with conventional read alignment and variant calling tools (see Materials and Methods). Allele balance plots of SNPs called from reads generated from synthetic diploids and polyploids were able to reproduce expected allele frequency plots at 100x for synthetic ploidies up to and including 8n (Figure 2A, Supplementary Figure 2). Synthetic diploids could be discerned from synthetic polyploids at lower coverages, but resolution was reduced below 50x. For read sets with five or more synthetic genotypes (≥ 5n), 50x coverage was not adequate to resolve individual peaks in the allele balance histograms, but plots produced were still inconsistent with the expectations of diploidy. At 10x coverage, it was not possible to visually discern diploids from polyploids by comparing the allele balance histograms (Figure 2A, Supplementary Figure 2). When the variance of allele balance for these same SNP calls was calculated, diploids could be differentiated from polyploids at all coverages from 10x to 100x (Figure 2B). For synthetic polyploids, variance of allele balance dropped at lower coverages; this was not observed in statistical simulations (Figure 1B, 2B). This reduced variance of allele balance could be due to the read alignment or variant calling stages. However, the variance of allele balance at 10x coverage was able to discriminate between the synthetic diploid and polyploid individuals using conventional software.

**Figure 2.**
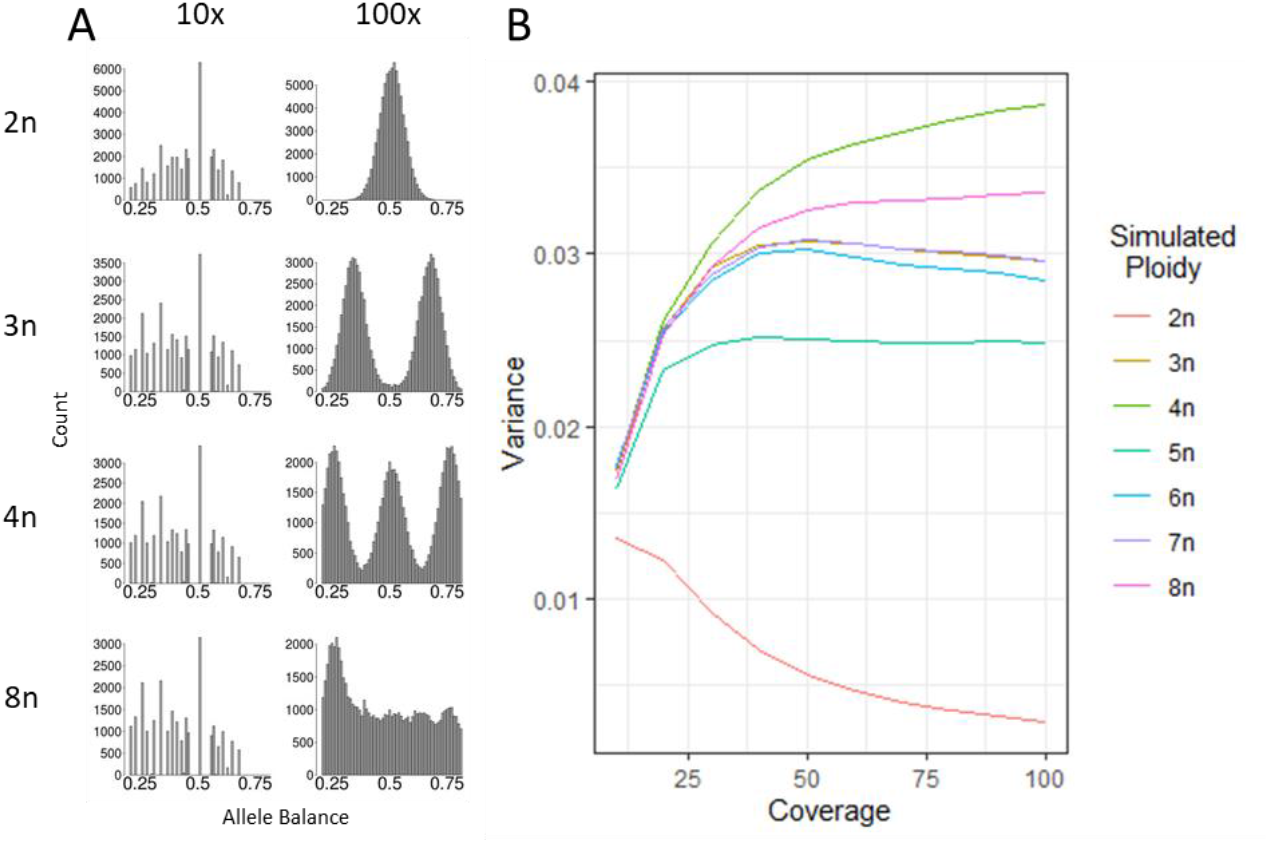
Variance of allele balance derived from reads generated for synthetic *E. coli* “polyploids”. A) Allele balance histograms produced from variant calls of synthetic *E. coli* datasets using simulated reads. At 10x it is hard to visually distinguish between diploids and polyploids due to the dominant 0.5x bar. At 100x it is possible to visually distinguish between ploidies. Additional downsampled allele balance histograms and ploidies are presented in Supplementary Figure 2. B) Variance of allele frequencies were calculated from the same data and could distinguish the synthetic diploid from synthetic polyploids. The variance of allele frequencies begins to converge at 10x, although they can still be differentiated.

When applied to real data, variance of allele balance was able to discriminate between homokaryotic and heterokaryotic isolates of *B. lactucae*. More SNPs were called in heterokaryotic isolates than homokaryotic isolates (Supplementary Figure 3). At high coverage, allele balance histograms were able to discern homokaryotic isolates from heterokaryotic isolates (Figure 3A). This is because homokaryotic isolates contain two genotypes, while heterokaryotic isolates contain four or more genotypes. When downsampled to lower coverages, it was not possible to distinguish homokaryons from heterokaryons using allele balance histograms (Figure 3A, Supplementary Figure 3). In contrast, at all coverage levels, the variance of allele balance was lower for the 12 homokaryotic isolates than for the 18 heterokaryotic isolates. Sampling these isolates at multiple coverages produced curves similar to that of the *E. coli* simulated data (Figure 2B, 3B). The isolates 1689 and 1806 displayed allele balance variances, which, while inconsistent with diploidy, did not fall within the range of other heterokaryotic isolates. The allele balance histograms showed that these isolates have unresolved profiles as previously reported (Fletcher, et al., 2019). The conclusion from that study was that these isolates are complex heterokaryons, possibly caused by an uneven mixture of multiple genotypes. Therefore, VCFvariance.pl is able to recapitulate previous conclusions (Fletcher, et al., 2019) based on lower sequencing coverage.

**Figure 3.**
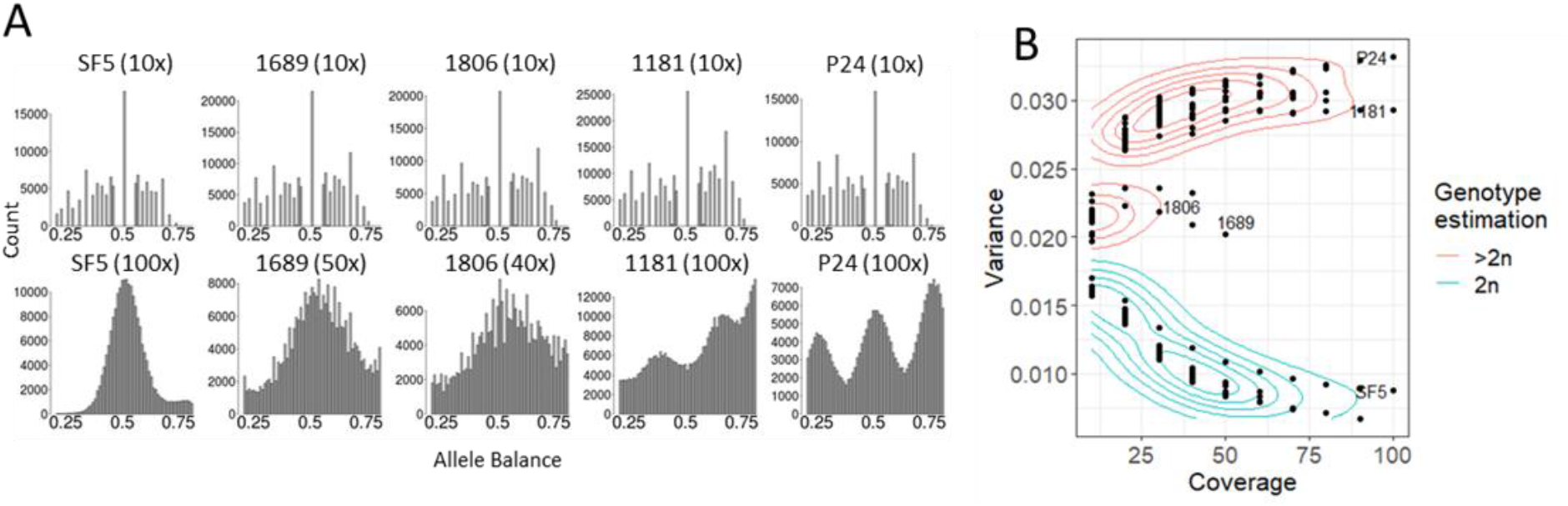
Variance of allele balance calculated for isolates of *B. lactucae* previously confirmed as homokaryotic or heterokaryotic. Whole-genome sequencing datasets of thirty isolates that had been sequenced to high coverage were downsampled to 10x to the highest coverage of factor by tens up to 100x. A) Allele frequency plots of isolates previously identified as homokaryotic (SF5) or heterokaryotic (all others). At 10x coverage it is not possible to distinguish the homokaryotic isolate from the heterokaryons. At higher coverage, multiple allele peaks are evident corresponding to different numbers of genotypes. B) Variance of allele balance was calculated from the same downsampled datasets. At 10x, homokaryotic and heterokaryotic isolates could be differentiated from one another. The difference in variance between the two groups increased as sequencing coverage increased. Smoothing through kernel density estimation was used to identify contours around the datasets. Labels indicate isolates presented in A. See the Results section for discussion of isolates 1689 and 1806 that appear different compared to other heterokaryotic isolates, but do not fit the expectations for diploidy at any coverage.

The protocol was further validated by downsampling high coverage sequencing data of 24 isolates of *S. cerevisiae*. This analysis surveyed nine diploids and eleven polyploids as well as four haploids. As expected, the variance of allele balance was higher for polyploids than for diploids (Figure 4). When downsampled to 10x, the variance of allele balance for isolate GSY1034 provided weak support for diploidy because it was on the edge of the contours defined by *B. lactucae*. At higher coverage, the isolate was clearly diploid. This demonstrates potential ambiguity at low coverage that may require additional sequencing to resolve. Histograms of allele balance were investigated for two high coverage datasets that, at high coverage, fell outside of the contours defined by *B. lactucae*. For isolate YJM954, the frequency of allele balance was bimodal, inconsistent with diploidy (Figure 4B). For YJM676, the frequency of allele balance was consistent with diploidy, but heavy-tailed (Figure 4C), resulting in slightly increased variance. Across all isolates, more SNPs were called and passed the allele balance filter for polyploid isolates than diploid isolates (Supplementary Figure 4), as found for *B. lactucae*. Running the four haploid isolates of *S. cerevisiae* through VCFvariance.pl demonstrated that haploids could be identified, because polymorphisms called versus the reference assembly should not be heterozygous in haploid isolates. For haploid isolates at 50x, between 161 to 190 SNPs were identified, of which only 4.2% to 19.3% passed the allele balance filter (see Methods); in comparison, at least 57% of the SNPs passed the allele balance filter at 50x coverage of diploids and polyploids. At 10x, between 135 to 231 SNPs were identified in haploids, of which 25% to 53.1% passed the allele balance filter; in contrast, at least 79% of the SNPs passed the allele balance filter at 10x coverage in diploids and polyploids. Consequently, the percentage of polymorphisms passing the allele balance filter could be used to distinguish haploids (Figure 4D, Supplementary Figure 4). Therefore, variance of allele balance can be used to discriminate between haploids, diploids, and polyploids in isolates of *S. cerevisae*.

**Figure 4.**
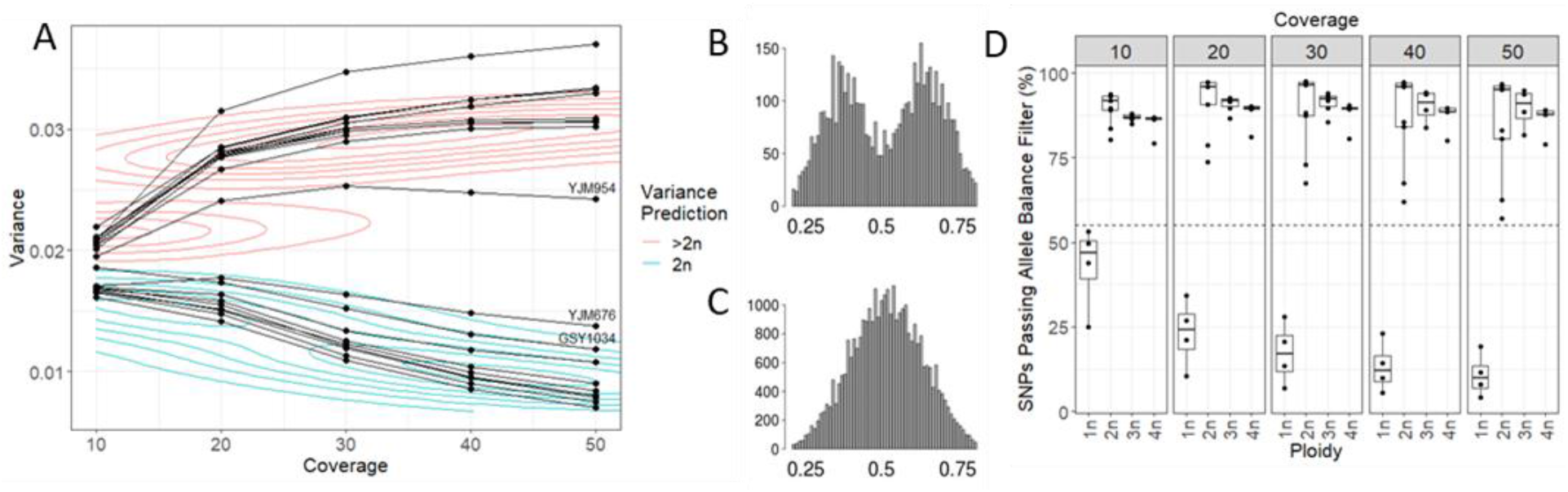
Variance of allele balance calculated for downsampled *S. cerevisiae* isolates exhibiting ploidy variation. Whole-genome sequencing datasets of isolates were downsampled to 10x, 20x, 30x, 40x, and 50x. A) Variance of allele balance was calculated from downsampled variant calls. Observations calculated from the same isolate are joined by a line. The contours provided as a guide were calculated using *B. lactucae* data (Figure 3). Histograms of allele balance were investigated for the two labelled isolates. B) Isolate YJM954 had bimodal distribution of allele balance frequencies inconsistent with diploidy. C) Isolate YJM676 had a heavy-tailed Gaussian distribution. D) The percentage of high-quality SNPs called as heterozygous could differentiate haploid isolates from polyploidy isolates. At all coverages the percentage of SNPs passing the allele balance filter never exceeded 55% for haploid isolates and was never less than 55% for diploid or polyploid isolates. The closest points were 53.1% for haploid isolate CBS1227 calculated from 135 polymorphisms at 10x and 57.1% for diploid isolate CBS2910 calculated from 2,377 SNPs at 50x. See Supplementary Figure 5 for more details.

Variance of allele balance was investigated for populations of *A. arenosa* that had been previously established as diploid or polyploid (Figure 5). The sequencing coverage of samples surveyed ranged from 10x to 35x. As with previous observations, more SNPs were identified for polyploids than for diploids and more SNPs were identified in individuals sequenced to higher coverage (Supplementary Figure 5). The variance of allele balance was higher for polyploids than for diploids and fell within the contours defined by heterokaryotic and homokaryotic isolates of *B. lactucae*. Therefore, variance of allele balance calculated for *A. arenosa* was able to discriminate between populations previously established to be diploid or polyploid.

**Figure 5.**
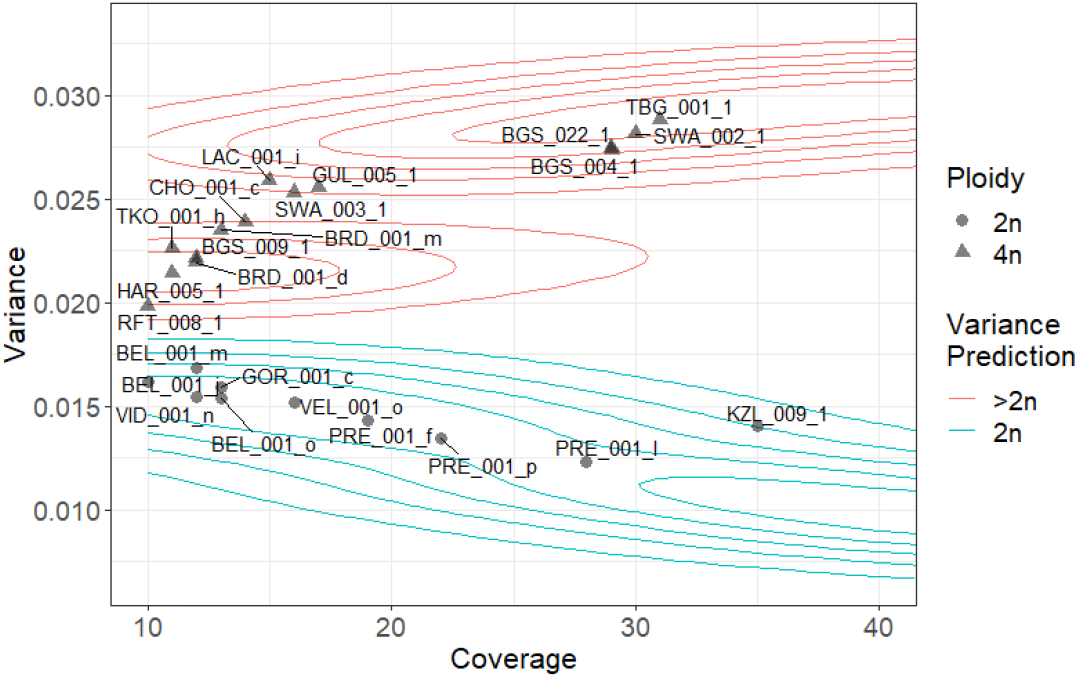
Variance of allele balance calculated for 24 *A. arenosa* individuals. Individuals previously described as diploid (circles) had a lower variance of allele balance compared to polyploid (triangles) individuals. Variance values calculated for all individuals fell within the contours defined for *B. lactucae* (Figure 3).

Variance of allele balance analysis for 40 *P. infestans* isolates identified 21 as diploid; 19 were inconsistent with diploidy (Figure 4). Most isolates fall within the contours defined by *B. lactucae*. These Consideration of the variance plots by geographic region demonstrated that all but one of the 16 isolates analyzed from Mexico were diploid, consistent with sexual reproduction occurring in the region. Four of the ten isolates surveyed from the USA, one of the seven from South America, and one of the six isolates surveyed from Europe were diploid. This panel included three herbarium specimens collected in the 1950s, labeled kew122, kew123, and kew126, that were all inconsistent with diploidy. These results are consistent with the previously reported polyploidy and copy number variation described for these isolates (Knaus, et al., 2019; Li, et al., 2016; Yoshida, et al., 2013).

Finally, VCFvariance.pl was deployed to investigate the nuclear state of type isolates for European races of *B. lactucae*. Sequences were generated for 13 isolates and variance of allele balance was calculated for these and eight previously reported isolates (Fletcher, et al., 2019) sequenced to high coverage and used to validate the protocol (Supplementary Tables 2 and 7). Twelve of the 21 isolates were diploid. The remaining nine, BL-EU01, BL-EU04, BL-EU05, BL-EU14, BL-EU19, BL-EU20, BL-EU30, BL-EU31, and BL-EU34, were inconsistent with diploidy and may therefore be heterokaryotic (Figure 7). The heterokaryotic nature of these isolates could result in unstable virulence phenotypes due to fluctuations in the populations of their nuclei (Fletcher, et al., 2019).

## Discussion

We show that the variance of allele balance is an effective measure to distinguish a sample with two genotypes (diploid, homokaryotic), from those with more than two genotypes (polyploid, heterokaryotic, or mixed culture). This was demonstrated using simulated (Figure 1), synthesized (Figure 2), and authentic sequencing data (Figure 3 to 6). This approach is robust and can be applied in multiple biological situations. In species with variable ploidy levels, such as *A. arenosa* (Monnahan, et al., 2019) and *S. cerevisae* (Zhu, et al., 2016), the variance of allele balance was consistently higher for polyploids compared to diploids at all coverages tested over or equal to 10x. For *B. lactucae*, it correctly differentiated previously characterized heterokaryotic isolates from homokaryotic isolates (Fletcher, et al., 2019) at all coverages tested and determined whether 13 race type isolates were homokaryotic. For *P. infestans*, it classified isolates as consistent or inconsistent with diploidy as previously established (Knaus, et al., 2019). This approach was also successfully applied to herbarium samples of *P. infestans* (Knaus, et al., 2019; Martin, et al., 2014). The variance of allele balance reflects the number of haplotypes present regardless of whether it originates from a plant, animal, or microbe. This meant that the same kernel density estimate contours could be plotted as a visual guide across all datasets (Figures 3-7). Therefore, calculating the variance of allele balance is a robust method for detecting deviations from the diploid state.

**Figure 6.**
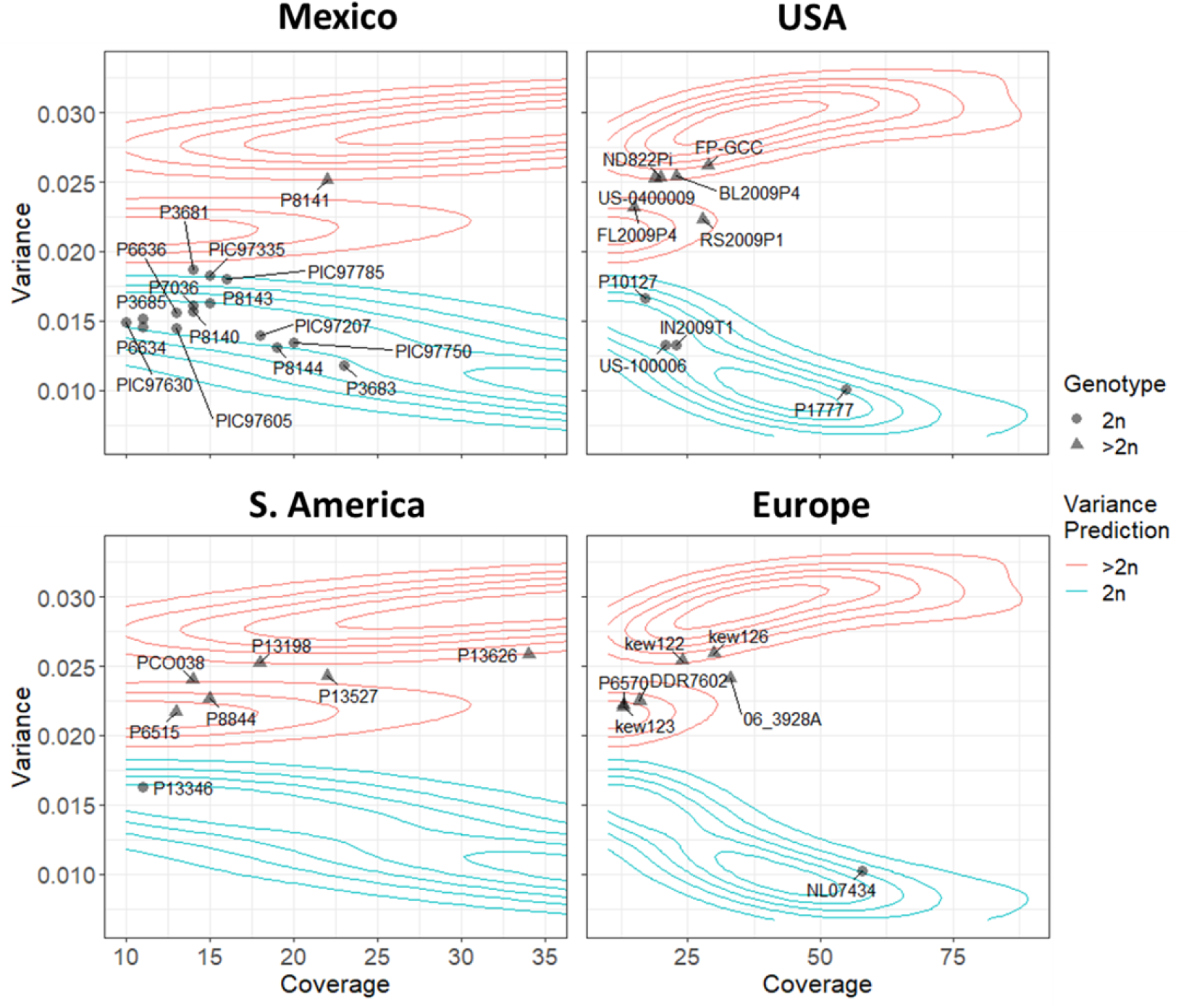
Variance of allele balance calculated for 40 isolates of *P. infestans*. Panels represent the geographic location the isolate was collected from as reported in other studies (Knaus, et al., 2019; Martin, et al., 2014). Isolates previously described inconsistent with diploidy (triangles) had higher variance of allele balance than isolates consistent with diploidy (circles). Variance values calculated for most isolates fell within the contours defined for *B. lactucae* (Figure 3).

**Figure 7.**
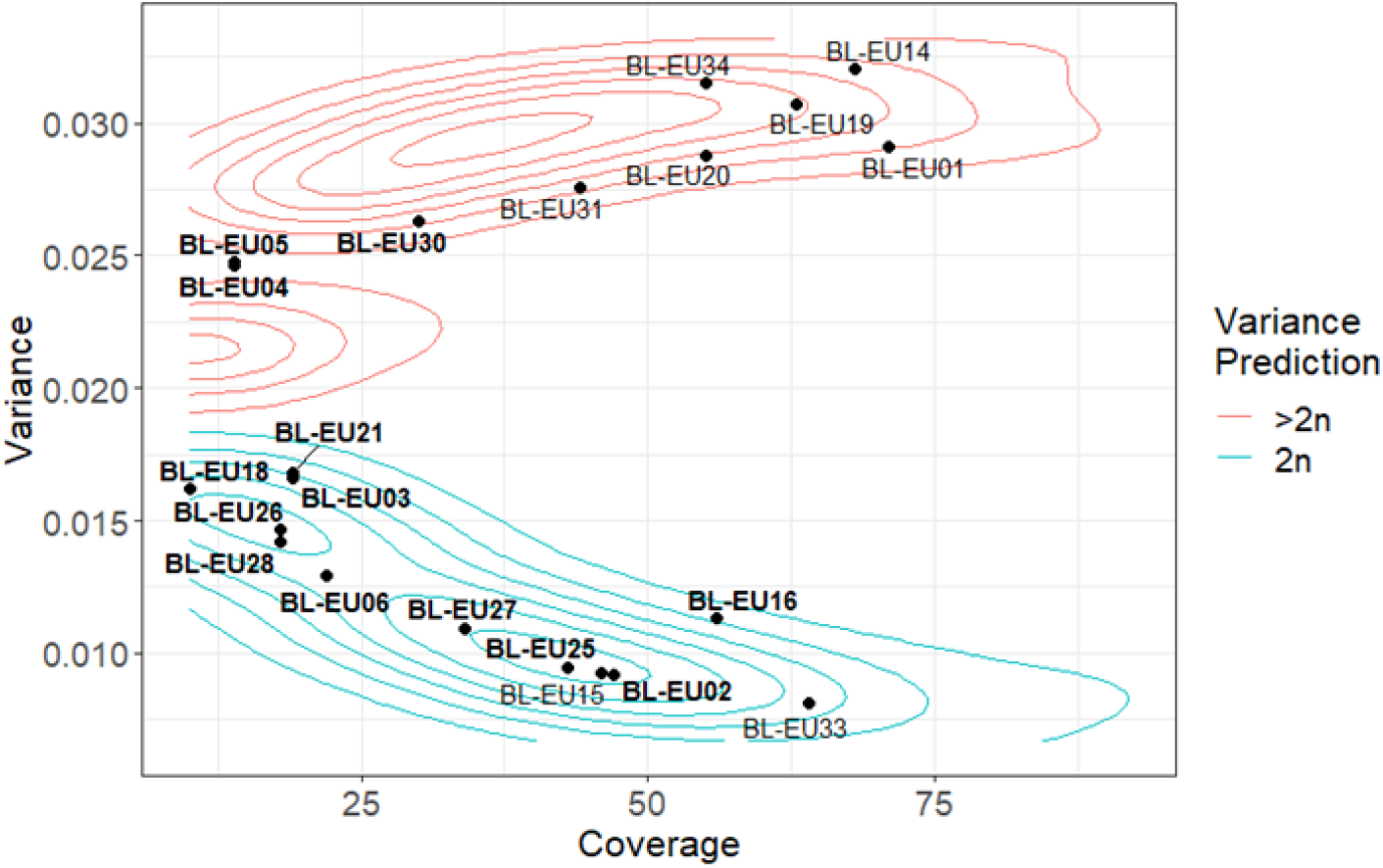
Variance of allele balance calculated for 21 European race type isolates of *B. lactucae*. Contours calculated in Figure 3 indicated that 12 isolates were homokaryotic. The other nine isolates are likely heterokaryotic and fall within contours calculated for heterokaryons. Bold-faced labels indicate 13 isolates for which the heterokaryotic condition was not previously known. Plain-faced labels indicate isolates for which histograms of allele balance have been reported (Fletcher, et al., 2019); these isolates were used to generate Figure 3.

This strategy is successful because an individual with more than two genotypes will have allele frequencies that significantly deviate from the expected 0.5/0.5 ratio (Fletcher, et al., 2019; Yoshida, et al., 2013). Calculating variance of allele balance enables interpretation at much lower coverages than the 50x used previously and can therefore reduce the cost of detecting polyploids and heterokaryons by up to 80% (sequencing to 10x rather than 50x). The shortcomings of plotting variance of allele balance are the same as plotting histograms of allele balance: such analyses may lead to false conclusions if there has been recent hybridization between two highly homozygous species or recent whole-genome duplication. In these instances, the expected allele balance of a tetraploid would be predominantly 1:1, despite there being four copies per locus. Variance of allele balance cannot delineate the number of genotypes in polyploids or heterokaryons at any coverage; manual inspection of allele balance plots generated from high coverage data retain value for determining the number of genotypes in a polyploid or heterokaryotic individual. Finally, at low coverages there may be ambiguity in the results because the variance of allele balance for diploids and polyploids begin to converge, as shown with downsampled simulated and authentic data (Figures 2-4). Data points that fall outside of the contours at low coverage may require higher coverage to conclude whether they are consistent with diploidy. Therefore, plotting the coverage on the x-axis conveys the confidence in the call made by VCFvariance.pl. An example of such a point can be seen in the *S. cerevisae* data, where at 10x the isolate GSY1034 borders the diploid call, but at higher coverages is confidently called to be diploid (Figure 4).

Calculating the variance of allele balance allows efficient and objective presentation of data while avoiding potential ambiguity. Presentation of allele balance histograms requires one plot per sample (Fletcher, et al., 2019; Knaus, et al., 2019); in contrast, communication of results from the variance of allele balance for multiple samples can be presented in a single plot. Interpretation of allele balance histograms can be subjective and may require experienced interpretation and explanation. Calculating the variance of allele balance provides a single value indicative of whether the genotype of an individual is consistent with diploidy. Therefore, using the variance of allele balance to infer departures from diploidy reduces ambiguity in the interpretation of results. This method provides a computationally light-weight solution that enables whole-genome ploidy analyses at low sequencing coverage.

## Conclusions

This study demonstrated a methodology for reliable detection of the departure from a diploid state using low coverage whole-genome sequencing data of any eukaryotic species. Such departures may be due to polyploidy, heterokaryosis, a mixed sample, or chromosomal copy number variation. This protocol can also detect haploid samples. Deployment of this strategy is computationally inexpensive and can reduce the sequencing costs by up to 80%. This protocol has been used to determine the heterokaryotic status of type isolates of *B. lactucae* and can be readily deployed to test for departures from diploidy in any organism.

## Contributions

KF conceptualized the project, performed the analysis, and drafted the manuscript. RH conceptualized the project and contributed to all drafts. DS provided biological material. RM supervised the project, contributed to data analysis, and all drafts. All authors contributed to writing and editing the paper and have approved the final submission.

## Acknowledgements

We thank K. Cavanaugh (UC Davis) for DNA extractions and preparation of Illumina paired-end libraries, H. Xu (UC Davis) for raw data submissions to NCBI, and E. Georgian (UC Davis) for editorial services. The sequencing was carried out by the DNA Technologies and Expression Analysis Cores at the UC Davis Genome Center, supported by NIH Shared Instrumentation Grant 1S10OD010786-01. The bioinformatic analysis was carried out using the UC Davis LSSC0 High Performance Computing cluster maintained by the UC Davis Bioinformatics Core.

## Data Availability Statement

Data downloaded from NCBI for validation of this approach are listed in Supplementary Tables 2 to 6. New data generated during this project are available under BioProject PRJNA387192.

---

**Supplementary Figure 1. Histograms of allele balance generated from simulated data**. The grid indicates expected allele balance for diploids through to octoploids sequenced from 10x to 100x.

**Supplementary Figure 2. Histograms of allele balance generated from synthetic *E. coli* data**. The grid indicates allele balances calculated from diploids through octoploids sequenced from 10x to 100x. Synthetic reads were processed using BWA-mem (Li, 2013) and Freebayes (Garrison and Marth, 2012). In contrast to Supplementary Figure 1, Supplementary Figure 2 demonstrates the difficulty in subjectively discerning diploids from polyploids at lower coverages.

**Supplementary Figure 3. Polymorphisms identified for 30 isolates of *B. lactucae*, downsampled to different coverages, used to generate Figure 3**. The outlined boxplots indicate the number of high-quality polymorphisms of *B. lactucae* and filled boxplots indicate the number polymorphisms that passed the allele balance filter (i.e., were heterozygous). The number of polymorphisms called for all isolates increased with coverage up to 50x. Beyond 50x the number of polymorphisms plateaued, indicating data saturation. At all coverages, more polymorphisms were identified for heterokaryons than homokaryons. A higher percentage of the polymorphisms passed the allele balance filter for homokaryons than heterokaryons, indicated by the red filled boxplots being closer to the red outlined boxplots.

**Supplementary Figure 4. Polymorphisms identified for 24 isolates of *S. cerevisiae*, downsampled to different coverages, used to generate Figure 4**. The outlined boxplots indicate the number of high-quality polymorphisms of *S. cerevisiae* and filled boxplots indicate the number of polymorphisms that passed the allele balance filter (i.e., were heterozygous). At all coverages, the number of polymorphisms identified for haploids was much lower than for diploids or polyploids. More polymorphisms were identified for tetraploids than triploids and for triploids than diploids. At all coverages, a higher percentage of the polymorphisms passed the allele balance filter for diploids (green) than triploids (turquoise) and for triploids than tetraploids (purple), indicated by the filled boxplots being closer to the outlined boxplots.

**Supplementary Figure 5. Polymorphisms analyzed for 24 *A. arenosa* individuals used to generate Figure 5**. The colors indicate whether the individual originates from a diploid or a polyploid population. Smaller circles indicating the number of high-quality polymorphisms are linked by a black line to larger circles, indicating the number of polymorphisms that passed the allele balance filter (i.e., were heterozygous). More polymorphisms were identified for individuals from polyploid populations than from diploid populations. The percentage of variants passing the allele balance filter was higher for individuals from diploid populations than for individuals from polyploid populations.

