## Supplementary figures and images for "Variance of allele balance calculated from low coverage sequencing data infers departure from a diploid state"

### Supplementary Figure 1

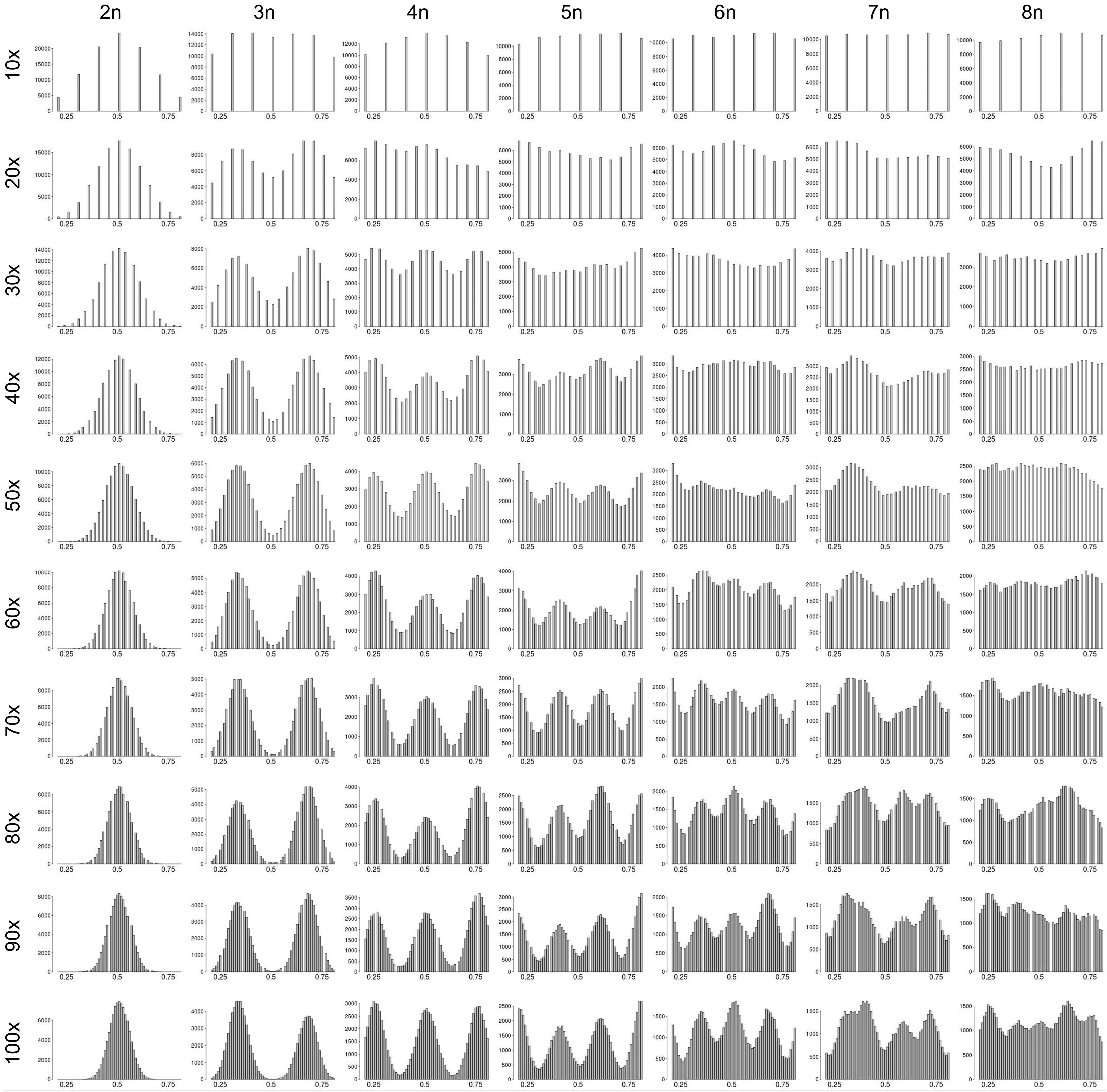

### Supplementary Figure 2

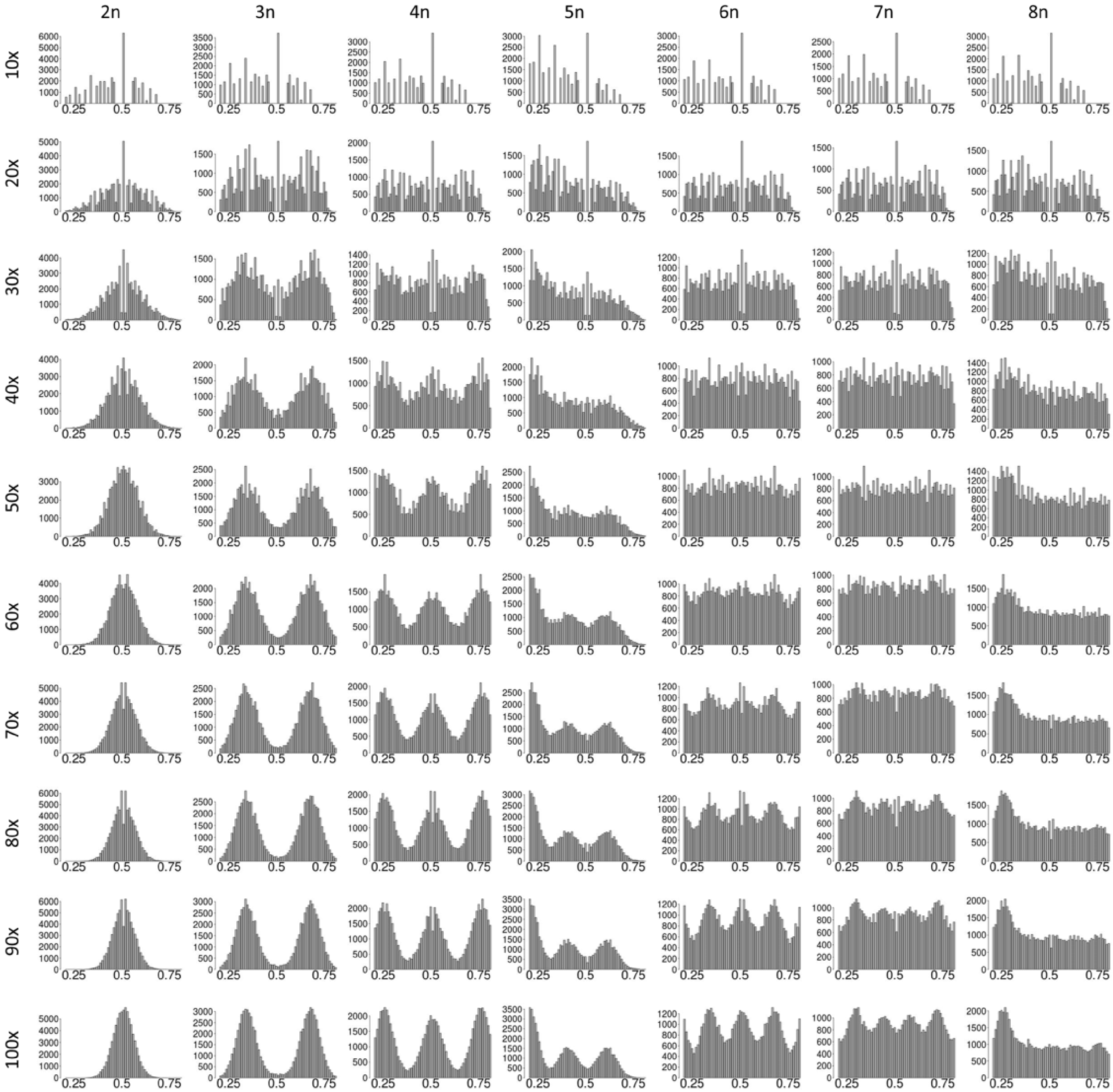
