## Supplementary Figures 3 to 5 for "Variance of allele balance calculated from low coverage sequencing data infers departure from a diploid state"

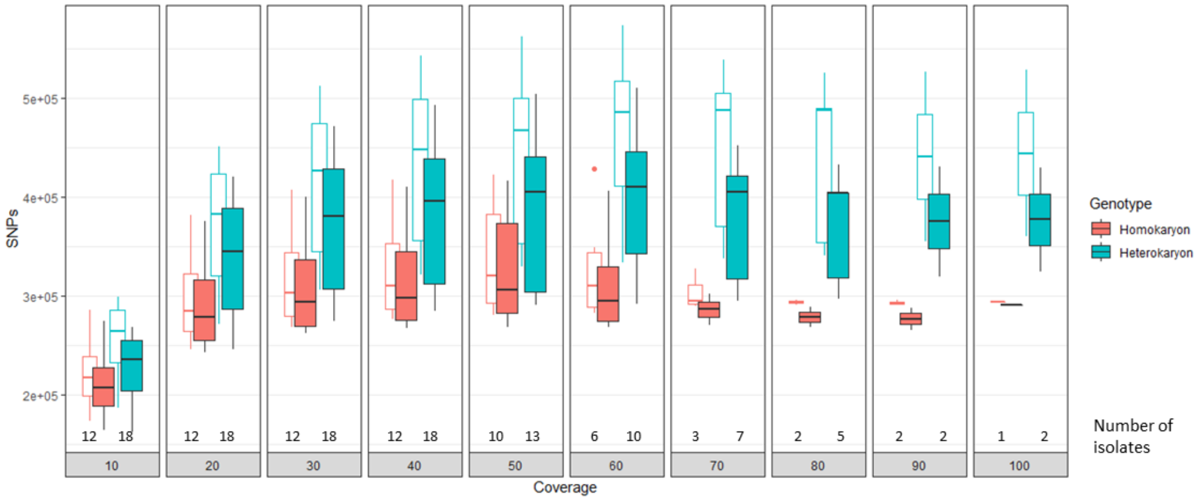


**Supplementary Figure 3**. **Polymorphisms identified for 30 isolates of *B. lactucae*, downsampled to different coverages, used to generate Figure 3.** The outlined boxplots indicate the number of high-quality polymorphisms of *B. lactucae* and filled boxplots indicate the number polymorphisms that passed the allele balance filter (i.e., were heterozygous). The number of polymorphisms called for all isolates increased with coverage up to 50x. Beyond 50x the number of polymorphisms plateaued, indicating data saturation. At all coverages, more polymorphisms were identified for heterokaryons than homokaryons. A higher percentage of the polymorphisms passed the allele balance filter for homokaryons than heterokaryons, indicated by the red filled boxplots being closer to the red outlined boxplots.

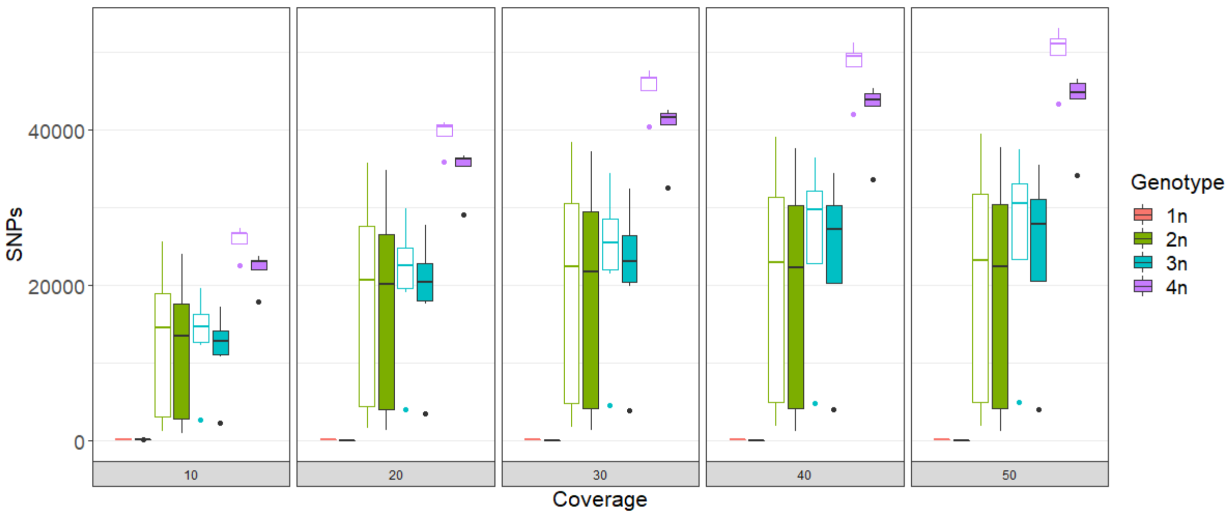


**Supplementary Figure 4. Polymorphisms identified for 24 isolates of *S. cerevisiae*, downsampled to different coverages, used to generate Figure 4.** The outlined boxplots indicate the number of high-quality polymorphisms of *S. cerevisiae* and filled boxplots indicate the number of polymorphisms that passed the allele balance filter (i.e., were heterozygous). At all coverages, the number of polymorphisms identified for haploids was much lower than for diploids or polyploids. More polymorphisms were identified for tetraploids than triploids and for triploids than diploids. At all coverages, a higher percentage of the polymorphisms passed the allele balance filter for diploids (green) than triploids (turquoise) and for triploids than tetraploids (purple), indicated by the filled boxplots being closer to the outlined boxplots.

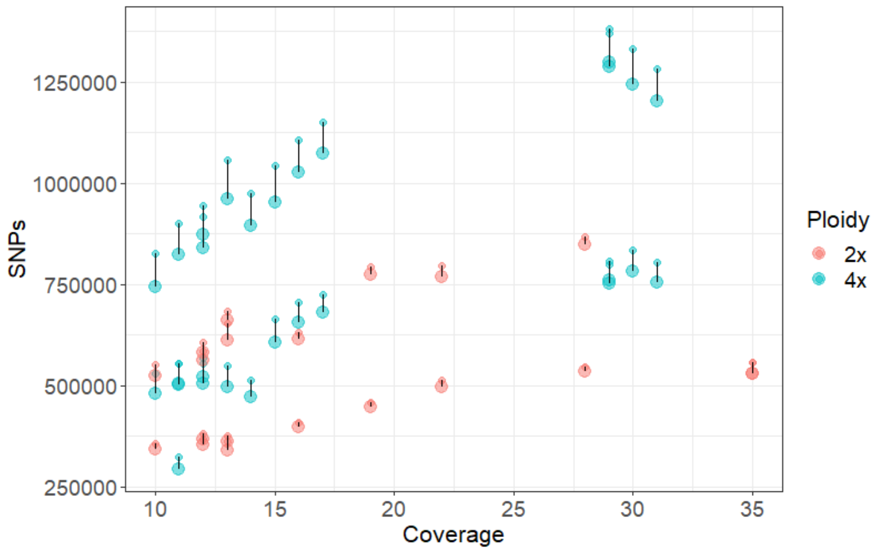


**Supplementary Figure 5. Polymorphisms analyzed for 24 *A. arenosa* individuals used to generate Figure 5.** The colors indicate whether the individual originates from a diploid or a polyploid population. Smaller circles indicating the number of high-quality polymorphisms are linked by a black line to larger circles, indicating the number of polymorphisms that passed the allele balance filter (i.e., were heterozygous). More polymorphisms were identified for individuals from polyploid populations than from diploid populations. The percentage of variants passing the allele balance filter was higher for individuals from diploid populations than for individuals from polyploid populations.
